# Prevalence of antimicrobial resistance phenotypes and genes in stable fly- and manure-derived bacterial isolates from clinically relevant taxa in dairy settings

**DOI:** 10.1101/2024.12.03.626706

**Authors:** Andrew J. Sommer, Julia E. Kettner, Travis K. Worley, Jordan Petrick, Caroline Haynie, Kerri L. Coon

**Author notes:** Corresponding author: Kerri L. Coon, 1550 Linden Drive, Madison, WI 53706.

## Abstract

**Aims:** This study aimed to characterize and compare the antimicrobial resistance (AMR) profiles of clinically relevant bacterial taxa isolated from biting stable flies (*Stomoxys* spp.) and bovine manure samples collected at a dairy research facility over the course of an entire fly breeding season. The presence of extended-spectrum beta-lactamase (ESBL) and other antimicrobial resistance genes (ARGs) was also examined.

**Methods and results:** A total of 606 fly- and 180 manure-derived strains were tested via disk diffusion for susceptibility to commonly administered antibiotics used in veterinary and human medicine. A small percentage of Enterobacterales exhibited resistance to the tested antimicrobials, including ceftiofur and other beta-lactam antibiotics. Extended spectrum beta-lactamase genes (TEM, CTX, OXA, CMY) were detected by PCR amplification in ceftiofur-resistant *Escherichia coli*, *Klebsiella* and *Enterobacter* spp. isolates. We additionally identified pirlimycin-resistant *Staphylococcus* and *Mammaliicoccus* spp. isolates encoding *lnuA*, a lincosamide resistance gene found primarily on small mobilizable plasmids.

**Conclusions:** These findings highlight the significance of stable flies in the carriage of antimicrobial-resistant bacterial strains and plasmid-associated ARGs on dairy farms.

## Impact statement

Livestock agricultural facilities represent important reservoirs of antimicrobial resistance. Our study confirms that biting *Stomoxys* stable flies can harbor multidrug resistant bacteria and may participate in the dissemination of ARGs on dairy farms. The carriage of ESBLs is of particular concern due to the use of cephalosporins in the treatment of bovine and human diseases.

## Introduction

The rise of antimicrobial resistance (AMR) represents a serious threat to global public health, causing an estimated five million deaths each year (Murray *et al*. 2022). In the United States, antimicrobial stewardship is of increasing concern to the dairy industry as agricultural microbiomes represent major reservoirs of both intrinsic and acquired antimicrobial resistance genes (ARGs) (Patel *et al*. 2020; Gelalcha and Kerro Dego 2022; Ruegg 2022). On dairy farms, the usage ceftiofur or other cephalosporin antibiotics remains a primary method of treatment for economically significant bovine diseases including mastitis, metritis, and enteritis (Sawant, Sordillo and Jayarao 2005; Redding, Bender and Baker 2019; Gonçalves *et al*. 2022). The continued usage of antimicrobials across agricultural and human healthcare settings has contributed to the acquisition, spread, and persistence of antimicrobial resistance in bacterial pathogens (Huttner *et al*. 2013; Ruegg 2022). AMR surveillance of dairy farms has focused primarily on bacterial strains isolated from bovine feces, clinical specimens, or food products (Makovec and Ruegg 2003; Oliver, Murinda and Jayarao 2011; Karp *et al*. 2017). Far less, however, is known about the carriage and potential maintenance of AMR by pest animals (flies, birds, rodents) coinhabiting the dairy barn ecosystem.

Stable flies (*Stomoxys* spp.) are ubiquitous and highly abundant pests of dairy farms (Hansens 1963; Meyer and Petersen 1983). Both male and female adult stable flies are obligate blood-feeders, which preferentially bite the underside or legs of dairy cattle (Dougherty *et al*. 1993; Showler and Osbrink 2015). Female flies will oviposit their eggs on aged or packed bovine manure, which serves as a protein, carbohydrate, and microbe rich source of nutrition for larvae (Meyer and Petersen 1983; Albuquerque and Zurek 2014). Our previous work has demonstrated that the microbiome of *Stomoxys* flies is environmentally derived, and that manure associated bacteria, including opportunistic bacterial pathogens, contribute significantly to the microbiome of adult flies captured from dairy farms (Sommer, Kettner and Coon 2024). Bacteria colonizing muscid flies may then be disseminated by adult flies through regurgitation, defecation, or on external surfaces, contributing to the potential environmental spread of bacteria both on and off farms (Butler *et al*. 1977; Sasaki, Kobayashi and Agui 2000; Baldacchino *et al*. 2013). Despite their status as significant pest species and their close association with important reservoirs of AMR such as manure, few studies have investigated the prevalence of antimicrobial-resistant bacteria in *Stomoxys* flies (Stelder *et al*. 2021; Gelalcha and Kerro Dego 2022).

In this study, we performed antimicrobial susceptibility testing on a collection of bacterial isolates derived from biting stable flies and environmental manure collected in parallel from Wisconsin dairies over a three-month period. A total of 786 bacterial isolates were tested for resistance to commonly administered human and veterinary antibiotics, providing one of the first comparative surveys of AMR rates in biting flies compared to manure. Notably, we identified flies as carriers of extended-spectrum beta-lactamase (ESBL)-encoding *Enterobacteriaceae* showing phenotypic resistance against ceftiofur and other cephalosporin antibiotics. We additionally identified pirlimycin-resistant *Staphylococcaceae*, which encoded the plasmid-borne *lnuA* (*linA*) lincosamide resistance gene. This study provides direct evidence to suggest that flies can act as carriers of drug-resistant opportunistic pathogens on dairy farms.

## Materials and methods

### Sample collection and bacterial isolations

Stable flies and manure were collected on a weekly basis from July through September 2021 at both the Emmons Blain Arlington Dairy Research Center (Arlington) and the UW-Madison Dairy Cattle Center (DCC), as previously reported (Sommer *et al*. 2024; Sommer, Kettner and Coon 2024). Briefly, flies were captured on adhesive fiberglass sticky traps (Olson Products, Medina, OH, USA), which selectively attract biting *Stomoxys* flies (Broce 1988). Biting flies were identified as *Stomoxys* with the assistance of taxonomic keys available in the Manual of Nearctic Diptera (MacAlpine 1993). A total of five traps were set and replaced weekly at the Arlington; a single trap was set outside the DCC. Retrieved trap liners were stored at -20 °C until further processing. Detailed methods for the generation of fly-bacterial homogenates and subsequent isolation of bacteria from endogenous and exogenous fly-surfaces are reported in two previous studies aimed at characterizing the taxonomic composition of the *Stomoxys* fly microbiota (Sommer *et al*. 2024; Sommer, Kettner and Coon 2024).

In this study, we performed further targeted isolations of *Staphylococcaceae*, *Klebsiella pneumoniae*, and *Escherichia coli* from bovine manure samples collected from the same dairy farms during the same timeframe the flies were captured. At least 25 ml of manure was collected from each of the five manure scraper systems at Arlington, which pool manure across the free-stall barn, as well as from both an isolated straw-bedding pen and from calf hutch areas. Manure from the DCC was collected from an outdoor pen area and from available fresh manure in the tie-stall barn stalls. Manure samples were diluted with sterile 1X phosphate-buffered saline (PBS) (1:1 ratio) and vortexed to ensure homogenization before storing at -80°C. Isolation of *E. coli* and *K. pneumoniae* was performed by streaking manure homogenates onto MacConkey agar (Hardy Diagnostics, Santa Maria, CA) and MacConkey-inositol-carbenicillin agar (base agar: HiMedia, Kenneth Square, PA; inositol: DOT Scientific, Burton, Mi; carbenicillin: Chem-Impex, Wood Dale, Il), which are selective for Gram-negative bacteria and *K. pneumoniae*, respectively (Bagley and Seidler 1978). Plates were incubated for 48 h at 37 °C. Presumptive *E. coli* colonies were confirmed as *E. coli* if a metallic green sheen was observed after streaking onto Eosin-Methylene Blue (EMB) agar (BD, Franklin Lakes, NJ) (Leininger, Roberson and Elvinger 2001). Presumptive *K. pneumoniae* colonies were streaked onto BHI and colony PCR was used to amplify a 1,400 bp region of the bacterial 16S rRNA gene using the universal primers 27F and 1492R (Frank *et al*. 2008). Amplicons were then sequenced via Sanger sequencing using the 1492R primer (Functional Biosciences, Madison, WI) and trimmed sequences were identified through BLAST comparison to the NCBI nr database (Altschul *et al*. 1990). Enrichment of Gram-positive bacteria was performed by streaking onto mannitol salt agar (Neogen, Lansing, MI) and incubating plates for 48 hours at 37° C. Potential *Staphylococcaceae* were identified using the *Staphylococcus* and *Mammaliicoccus* (previously classified as *Staphylococcus*) genus-specific primers TStaG422 and TStag765 (Martineau *et al*. 2001). Isolated strains were stored as 40% glycerol stocks at -80° C.

### Antimicrobial susceptibility testing of bacterial isolate collections

Antimicrobial susceptibility testing was performed via the disk diffusion method in accordance with Clinical and Laboratory Standards Institute (CSLI) guidelines. Testing of Enterobacterales isolates was performed with ampicillin (10 µg), chloramphenicol (30 µl), ceftiofur (30 µl), cephalothin (30 µg), gentamicin (30 µl), and tetracycline (30 µl). Enterobacterales that exhibited resistant or intermediate phenotypes against ceftiofur were further tested with ceftazidime (30 µg), ceftazidime (30 µg) + avibactam (20 µg), imipenem (10 µg), meropenem (10 µg), ertapenem (10 µg), and levofloxacin (5 µg). Testing of *Staphylococcaceae* isolates was performed with cefoxitin (30 µl), chloramphenicol (30 µl), ceftiofur (30 µl), gentamicin (10 µl), and pirlimycin (2 µl). Testing of *Enterococcus* isolates were performed with ampicillin (10 µl), chloramphenicol (30 µl), penicillin (10 units), and tetracycline (30 µl). Testing of *Acinetobacter* isolates was performed with gentamycin (10 µl), tetracycline (30 µl), and ceftazidime (30 µl). Breakpoints were selected based on recommendations in the CLSI M100 ED34 (CLSI 2024a), with the exceptions of CLSI VET01S ED7 (CLSI 2024b) for pirlimycin and ceftiofur, and M- 100-S25 for cephalothin (CLSI 2015). *E. coli* ATCC 25922 and *S. aureus* ATCC 25923 were used as control strains to validate testing methods.

### PCR detection of antimicrobial resistance genes

Colony PCR was used to determine the carriage of specific ARGs by strains exhibiting a resistant or intermediate phenotype to either ceftiofur or pirlimycin. Universal primers were used to test Enterobacterales isolates for the presence of the following extended-spectrum beta-lactamases (ESBLs): CTX, TEM, OXA-1, and CMY (Zhao *et al*. 2001; Pagani *et al*. 2003; Weill *et al*. 2004; Ogutu *et al*. 2015). *Staphylococcaceae* isolates were tested for the presence of the *lnuA* (*linA*) plasmid-borne lincosamide resistance gene (Wendlandt *et al*. 2015). Amplifications were performed as 25 µl reactions under the following conditions: initial denaturation cycle of 95°C for 5 min, followed by 30 cycles at 95°C for 15 s, annealing for 15 s, 68°C for 30 s, and a final extension step at 72°C for 5 min. Primer sequences and annealing temperatures are listed in Table S1. PCR products were visualized via gel electrophoresis on 1% agarose gels.

### Conjugation of bla-CTX

*E. coli* K-12 derivative BW25113 was chosen as a recipient strain and was chromosomally tagged with a Tn7 transposon carrying genes for resistance to both kanamycin and chloramphenicol (Datsenko and Wanner 2000). This transposon was delivered from the pMRE- Tn7-152 vector (Schlechter *et al*. 2018) using a previously described protocol (McKenzie and Craig 2006). For the conjugation, cultures of the ceftiofur resistant donor AS0255 and the BW25113-Tn7-152 recipient were grown overnight in lysogeny broth (LB) media and then subcultured 1:100 in 10 ml of LB broth to late logarithmic phase, approximately 0.8 – 1.0 OD_600_ . Donor and recipient cultures were then mixed at a 1:1 ratio to a final volume of 1 ml in a 1.5 ml centrifuge tube, pelleted at 11,000 x *g* for 2 min, and resuspended in 25 µl of LB. Donor and recipient cells were prepared individually the same way to serve as negative controls. The conjugation reaction and controls were then spot plated on individual LB agar plates and incubated overnight at 37 °C. Finally, the grown cells were scraped into 1 ml 1X PBS, pelleted and resuspended in 1 ml fresh PBS, and serially diluted 10-fold in PBS to 10^-6^. Dilutions were then spot plated in duplicate onto LB agar plates containing either 30 µg/ml ceftiofur sodium, the combination of 50 µg/ml kanamycin and 15 µg/ml chloramphenicol, or all three antibiotics at the listed concentrations, and were incubated overnight at 37 °C. The number of transconjugants obtained was compared to the number of recipient cells to calculate the transfer efficiency (transconjugants/donors) of the conjugation.

## Results

### Summary of tested isolates

A bacterial isolate collection derived from biting fly and manure samples collected from Wisconsin dairies was screened for susceptibility against common antimicrobials used in human and veterinary medicine. The isolate collection contains a total of 606 fly- and 180 manure-derived strains identified as belonging to one of the following taxonomic groups: Enterobacterales, *Staphylococcaceae*, *Acinetobacter*, and *Enterococcus* (Tables S2-S7). The most common bacterial genera in the collection include *Enterobacter*, *Escherichia*, *Klebsiella*, *Pantoea*, and non-aureus staphylococci and mammaliicocci.

### Antimicrobial susceptibility of fly- and manure-derived Enterobacterales

Disk diffusion was used to test 66 fly-derived and 46 manure-derived *E. coli* isolates for susceptibility against ampicillin, chloramphenicol, ceftiofur, cephalothin, gentamicin, and tetracycline antibiotics (Figure 1A; Table S2). Resistance to ampicillin, cephalothin, and tetracycline was most common across tested isolates derived from both sources; however, resistant strains represented a minority of the collection. Cephalothin was the only antibiotic against which the majority of *E. coli* isolates displayed either a resistant or intermediate phenotype, which included 51.5% of fly- and 63.0% of manure-derived strains. Ceftiofur resistance was observed for only a single fly-derived *E. coli* isolate and no manure-derived *E*.

**Figure 1.**
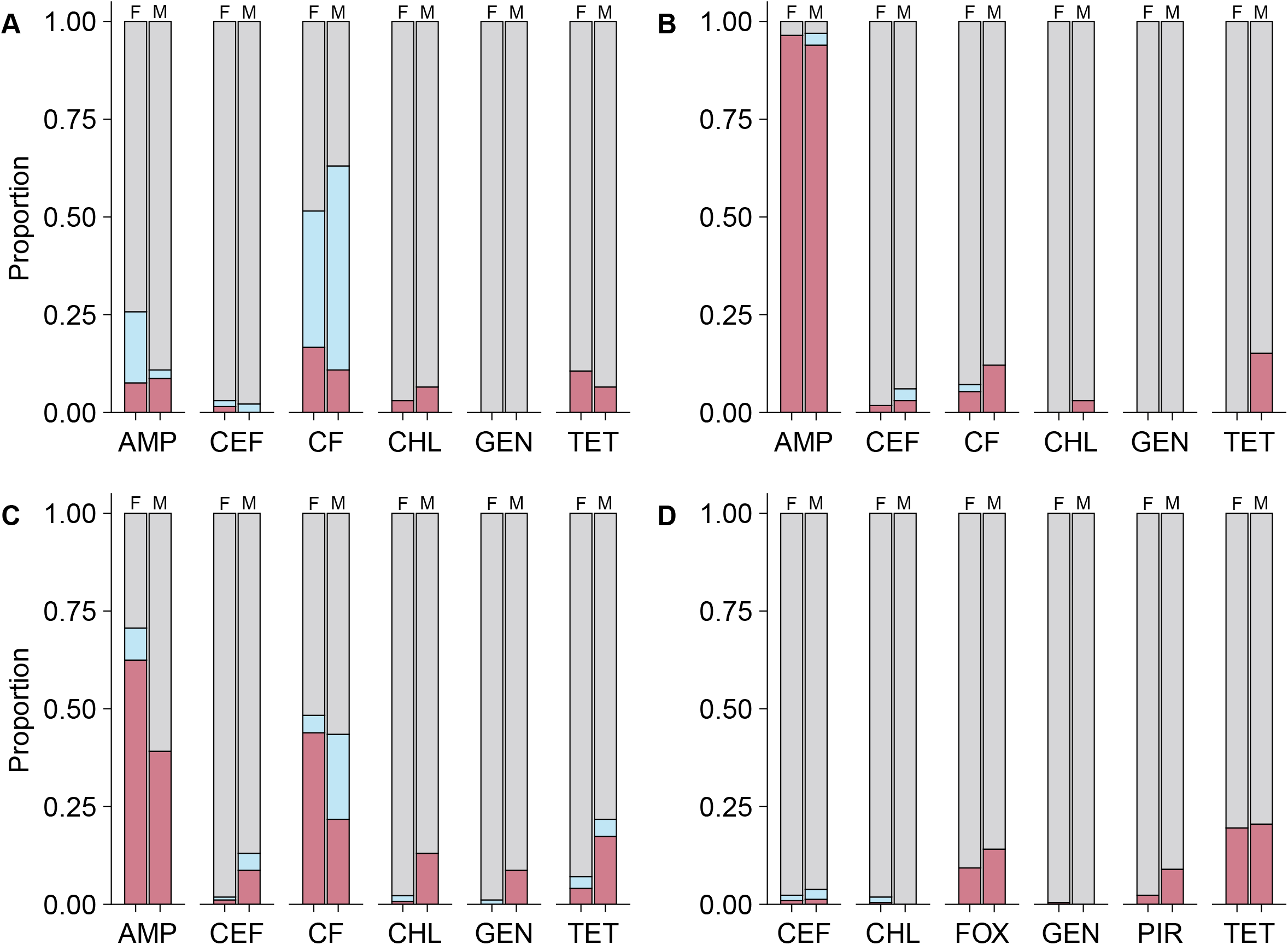
Antimicrobial susceptibility of Enterobacterales and Staphylococcaceae isolates. Paired bar charts show the proportion of isolates within each taxonomic group (A- *E. coli*, B- *Klebsiella*, C- Other Enterobacterales, D- Staphylococcaceae) that exhibited resistant (bottom/red), intermediate (middle/blue), or susceptible (top/grey) phenotypes against a given antibiotic. Paired bars marked with an “F” represent fly-derived isolates, while paired bars marked with an “M” represent manure-derived isolates. All Enterobacterales (Panels A-C) were tested against ampicillin (AMP), ceftiofur (CEF), cephalothin (CF), chloramphenicol (CHL), gentamycin (GEN), and tetracycline (TET). All Staphylococcaceae were tested against ceftiofur (CEF), chloramphenicol (CHL), cefoxitin (FOX), gentamycin (GEN), and tetracycline (TET).

*coli* isolates. Gentamicin resistance was not observed for any *E. coli* isolates. Resistance to ampicillin was highly prevalent across almost all of the 89 *Klebsiella* isolates tested (95.5%) (Figure 1B; Table S3). Contrary to *E. coli* strains, *Klebsiella* strains displayed low levels of resistance to cephalothin (7.9%). Ceftiofur resistance was observed for a single fly-derived and a single manure-derived isolate, while gentamicin resistance was not found in any *Klebsiella*. Among the other 292 Enterobacterales isolates, we found high levels of resistance to ampicillin in *Enterobacter* (84.6%), *Kosakonia* (95.7%), and *Serratia* spp. isolates (97.3%) (Figure 1C, Table S4). *Serratia* and *Enterobacter* spp. isolates also showed high levels of resistance to cephalothin (100% and 93.6%, respectively). Three *Enterobacter*, one *Pantoea*, and one *Proteus* exhibited resistance to ceftiofur.

All Enterobacterales with resistant or intermediate phenotypes for ceftiofur were subjected to an additional panel of susceptibility testing consisting of ceftazidime with and without the beta-lactamase inhibitor avibactam, imipenem, meropenem, ertapenem, and levofloxacin (Table S8). Almost all strains that were resistant to ceftiofur were also resistant to ceftazidime, another 3^rd^ generation cephalosporin. However, a ceftiofur-resistant *Proteus* isolate was susceptible to ceftazidime, while a *Klebsiella* and an *E. coli* that were intermediate for ceftiofur were resistant to ceftazidime. All strains except for the ceftiofur-resistant *Pantoea* isolate were susceptible to the combination of ceftazidime and the beta-lactamase inhibitor avibactam. Notably, this same *Pantoea* isolate was also resistant to meropenem and intermediate for imipenem and ertapenem, all three of which are carbapenems. Beyond this strain, carbapenem (ertapenem) resistance was detected in one *Kosakonia* isolate and two Enterobacter isolates displayed an intermediate phenotype to ertapenem. All strains were susceptible to the fluoroquinolone levofloxacin.

### Antimicrobial susceptibility of fly- and manure-derived Staphylococcaceae

A total of 215 fly- and 78 manure-derived *Staphylococcaceae* isolates were tested via disk diffusion for susceptibility against cefoxitin, chloramphenicol, ceftiofur, gentamicin, and pirlimycin (Figure 1D; Table S5). Fly- and manure-derived isolates displayed the highest prevalence of resistance to tetracycline compared to other tested antibiotics, with 19.5% and 20.5% of the isolates showing a resistant phenotype, respectively. Conversely, only a single isolate exhibited a resistant phenotype to either chloramphenicol or gentamicin. We found a total of two fly-derived isolates and one manure-derived isolate that showed resistance to ceftiofur. Similarly, five fly-derived and seven manure-derived isolates showed resistance to pirlimycin.

### Antimicrobial susceptibility of Acinetobacter and Enterococcus spp. isolates

A total of 45 *Acinetobacter* strains and 10 *Enterococcus* strains, primarily obtained as by- products of other isolations, were also tested using disk diffusion. We found that none of the tested *Acinetobacter* strains showed resistance to gentamicin, tetracycline, or ceftazidime (Table S6). Similarly, no *Enterococcus* strains showed resistance to ampicillin, chloramphenicol, or penicillin (Table S7).

### ARG carriage by ceftiofur-resistant Enterobacterales and pirlimycin-resistant Staphylococcaceae

Enterobacterales strains with either an intermediate or resistant ceftiofur phenotype were further screened for the presence of select ESBL genes to determine potential mechanisms of ceftiofur resistance in the tested isolates. Notably, a subset of these strains was also resistant to non-beta-lactams, including tetracycline and gentamycin (Table 1). ESBLs were detected in five of the nine ceftiofur-resistant isolates (Table 1), which showed cross resistance to ampicillin and cephalothin antibiotics. While bla-CTX was observed exclusively in ceftiofur-resistant strains, bla-CMY was detected in both a *Klebsiella* (AS0507) and *E. coli* (AS0540) strain that showed an intermediate phenotype to ceftiofur (File S2). In contrast, bla-OXA was observed in only a single strain (*Enterobacter* AS0467), in which it was encoded alongside a CTX and TEM gene. We also identified one strain (*E. coli* AS0466) encoding bla-TEM, but not bla-OXA or bla-CTX, which did not exhibit resistance to ceftiofur (File S2). Follow-up conjugation experiments demonstrated that ceftiofur resistance could be transferred from a bla-CTX-encoding strain (*E. coli* AS0255) to BW25113, an *E. coli* K-12 derivative, at an average rate of 1.36 x 10^-5^ transconjugants/recipient, confirming that encoded ESBL genes and associated resistance to cephalosporins exhibited by some of the strains we tested are likely plasmid-borne (Table S9).

**Table 1.** Antimicrobial resistance profiles and detection of extended-spectrum beta-lactamases (ESBLs) in ceftiofur-resistant Enterobacterales.

| Strain | Taxon | Source | Resistance profile | ESBL genes detected |
| --- | --- | --- | --- | --- |
| AS0467 | *Enterobacter* | Manure | AMP, CEF, CF, CHL, GEN, TET | CTX, TEM, OXA |
| AS1210 | *Enterobacter* | Fly | AMP, CEF, CF | None |
| AS1211 | *Enterobacter* | Fly | AMP, CEF, CF | None |
| AS0255 | *Escherichia* | Fly | AMP, CEF, CF, TET | CTX, TEM |
| AS0468 | *Klebsiella* | Manure | AMP, CEF, CF, CHL, TET | CTX |
| AS1245 | *Klebsiella* | Fly | AMP, CEF, CF | CTX |
| AS0690 | *Pantoea* | Fly | AMP, CEF, CF | None |
| AS0234 | *Proteus* | Manure | CHL, CEF, CF, GEN, TET* | None |

\*Indicates an intermediate phenotype was observed.
Abbreviations: ampicillin (AMP), ceftiofur (CEF), cephalothin (CF), chloramphenicol (CHL), gentamycin (GEN), and tetracycline (TET).

Finally, pirlimycin-resistant *Staphylococcaceae* were tested for the presence of the *lnuA* lincosamide resistance gene. We detected *lnuA* in 11 of the 12 pirlimycin-resistant strains (Table 2), while the *lnuA* gene was not detected in any of the seven pirlimycin-susceptible strains tested for comparison (File S2).

**Table 2.** Antimicrobial resistance profiles and detection of *lnuA* in pirlimycin-resistant *Staphylococcaceae*.

| Strain | Taxon | Source | Resistance profile | *lnuA* detected |
| --- | --- | --- | --- | --- |
| AS0295 | *Mammaliicoccus* | Fly | PIR | Yes |
| AS0311 | *Staphylococcus* | Manure | PIR | Yes |
| AS0554 | *Mammaliicoccus* | Manure | FOX, CEF, PIR | Yes |
| AS0569 | *Staphylococcus* | Manure | PIR | Yes |
| AS0602 | *Staphylococcus* | Fly | PIR, TET | Yes |
| AS0913 | *Mammaliicoccus* | Fly | PIR, TET | Yes |
| AS1275 | *Staphylococcus* | Fly | PIR, TET | Yes |
| AS1301 | *Staphylococcus* | Fly | PIR, TET | Yes |
| AS1397 | *Staphylococcaceae* | Manure | PIR | No |
| AS1400 | *Mammaliicoccus* | Manure | PIR, TET | Yes |
| AS1404 | *Staphylococcaceae* | Manure | PIR, TET | Yes |
| AS1445 | *Staphylococcus* | Manure | PIR, TET | Yes |

Abbreviations: ceftiofur (CEF), chloramphenicol (CHL), cefoxitin (FOX), gentamycin (GEN), and tetracycline (TET).

## Discussion

The continued usage of antibiotics has contributed to the spread of antimicrobial resistance and treatment-resistant pathogens in both animal and human medicine. A “One Health” perspective, which recognizes the interconnectedness of humans, animals, and the environment, is essential to understand the spread and dispersal of antimicrobial resistance across ecosystems. Despite their suspected carriage of opportunistic pathogens and close association with manure reservoirs on dairy farms (Baldacchino *et al*. 2013; Sommer, Kettner and Coon 2024), biting *Stomoxys* stable flies represent understudied elements of antimicrobial resistance transmission. Here, we performed antimicrobial susceptibility testing on a collection of 786 bacterial isolates derived from *Stomoxys* flies and nearby manure piles. Our results reveal that *Stomoxys* flies can serve as carriers of multidrug resistant bacterial strains, which emphasizes the need to expand antimicrobial resistance surveillance to include flies on dairy farms.

In this study, we determined the rates of beta-lactam antibiotic susceptibility and detected the potential mechanisms of ceftiofur resistance in a large collection of bacterial strains isolated from dairy farms. Ceftiofur and other third-generation cephalosporins are the most widely administered antimicrobials for the treatment of bovine mastitis and other economically significant diseases on dairy farms in the United States (Sawant, Sordillo and Jayarao 2005; Redding, Bender and Baker 2019; Gonçalves *et al*. 2022). Cephalosporin antibiotics are also routinely used in the treatment of human diseases as they provide broad-spectrum coverage against both Gram-positive and Gram-negative bacterial pathogens (Lin and Kück 2022). Resistance to third-generation cephalosporins in *Enterobacteriaceae* is often mediated through the production of ESBLs, which also provide cross-resistance to other beta-lactams (ampicillin, penicillin, etc.) and are often encoded as mobile genetic elements on plasmids (Castanheira, Simner and Bradford 2021; Lin and Kück 2022). In this study, we found both fly- and manure-derived *E. coli*, *Klebsiella* and *Enterobacter* spp. strains exhibiting phenotypic resistance to ampicillin, cephalothin, and ceftiofur encoding a combination of the CTX, TEM, and OXA beta-lactamases. We also identified one *Klebsiella* and one *Escherichia* isolate displaying an intermediate level of resistance to ceftiofur, which was potentially mediated through the CMY beta-lactamase. We found that a bla-CTX-encoding *E. coli* isolate was able to transfer ceftiofur resistance to a naïve recipient strain of *E. coli* (Table S8), which indicates that the CTX gene is likely present on a conjugative plasmid in this strain. The dissemination of ESBLs via conjugative plasmids is considered a major driver of the spread of resistance to all classes of beta-lactam antibiotics (Pfeifer, Cullik and Witte 2010). We were not able to identify a potential mechanism of ceftiofur resistance for two *Enterobacter* isolates, one *Proteus* isolate, and one *Pantoea* isolate, suggesting that these strains could encode other ESBLs or ARGs not tested in this study. However, this *Pantoea* isolate and a *Kosakonia* isolate showed resistance to meropenem and ertapenem, respectively, suggesting the potential presence of carbapenemase genes in these strains. Carbapenems are important antibiotics for human health, and the rise of carbapenem resistance is considered a significant emerging threat to the ability to treat bacterial infections (Pfeifer, Cullik and Witte 2010).

Non-aureus staphylococci and mammaliicocci are major causative agents of persistent subclinical mastitis and are considered potential reservoirs of antimicrobial resistance genes on dairy farms (Taponen and Pyorala 2009; Schoenfelder *et al*. 2017). Resistance to beta-lactam antibiotics in *Staphylococcaceae* is commonly attributed to chromosomal transposon-encoded *mecA* genes, which display a large degree of genetic variability (Becker, Heilmann and Peters 2014). Past studies have also reported discrepancies between phenotypic resistance to beta-lactam antibiotics and the presence of *mecA* genes (Goncalves *et al*. 2023). We found that a higher prevalence of *Staphylococcaceae* isolates showed resistance to cefoxitin (a proxy for methicillin) compared to ceftiofur, highlighting the complexity of beta-lactam resistance in *Staphylococcaceae*. Other studies have also reported lower levels of ceftiofur resistance compared to other beta-lactam antibiotics (ampicillin, penicillin) in *Staphylococcus* species (Kim *et al*. 2019; Kolar *et al*. 2024).

Pirlimycin is a lincosamide antibiotic used in the treatment of intramammary infections caused by *Staphylococcus* and other Gram-positive bacteria (Thornsberry *et al*. 1993; Sawant, Sordillo and Jayarao 2005; Gonçalves *et al*. 2022). Prior work has established that lincosamide and macrolide resistance in *Staphylococcus* can be mediated through the expression of ARGs carried on mobile genetic elements (Lüthje, Von Köckritz-Blickwede and Schwarz 2007; Schoenfelder *et al*. 2017; Feßler *et al*. 2018). In this study, we screened isolates for the presence of the *lnuA* gene, which is primarily found on small rolling circle replication plasmids (Lüthje, Von Köckritz-Blickwede and Schwarz 2007). We detected *lnuA* in all but one pirlimycin-resistant *Staphylococcus* and one *Mammaliicoccus* spp. isolate that we tested, suggesting that pirlimycin resistance was driven by a plasmid-borne *lnuA* gene. Our results do not preclude the possibility that these strains carry other ARGs (*erm*, *cfr*, *vga*, *sal*) that also contribute to pirlimycin resistance (Feßler *et al*. 2018).

Antimicrobial susceptibility testing was also performed for infrequent or atypical opportunistic Enterobacterales pathogens, including isolates of *Pantoea* and *Kosakonia*. We found that nearly all *Kosakonia* isolates tested showed resistance to ampicillin and susceptibility to cephalothin. Intrinsic resistance to ampicillin is likely driven by the KSA-1, a recently characterized chromosomally encoded beta-lactamase in *Kosakonia sacchari* (Fournier *et al*. 2024). *Pantoea* isolates were largely susceptible to all antibiotics tested; however, we found one strain exhibiting multi-drug resistance to ceftiofur, ampicillin, and cephalothin. We did not detect the presence of the CTX, TEM, CMY, or OXA beta-lactamase genes, suggesting a different mechanism of resistance in this strain. Further work will be needed to determine the role of *Pantoea* or other *Erwiniaceae* species in the carriage of ARGs on dairy farms.

Collectively, our results provide evidence that biting flies can serve as carriers of antimicrobial resistant bacteria within the dairy farm environment. Further studies to elucidate the role of flies in the dissemination of ESBLs and other plasmid-borne ARGs are necessary to understand how the fly microbiota is interconnected with other elements of the dairy farm ecosystem. These insights will contribute to our understanding of the spread and persistence of antimicrobial resistance within agricultural food systems.

## Acknowledgements

We thank Jessica Cederquist and the UW-Madison Department of Animal and Dairy Science Dairy Herd Operation for access to the dairy cattle facilities included in this study (Arlington and the DCC), along with their associated records. We also thank Garret Suen for helpful conversations regarding study design, and well as the UW-Madison Department of Bacteriology teaching laboratory for providing an antibiotic disk stamper.

## Author contributions

A.J.S. (Conceptualization, Methodology, Validation, Formal analysis, Investigation, Data Curation Writing - original draft, Writing – Review & Editing, Visualization, Supervision);

J.E.K. (Methodology, Investigation, Data Curation, Writing - Review & Editing); T.K.W. (Methodology, Investigation, Data Curation, Writing - Review & Editing), J.P. (Methodology, Investigation, Data Curation); C.H. (Methodology, Investigation, Data Curation); K.L.C. (Conceptualization, Methodology, Validation, Writing - Review & Editing, Supervision, Project Administration, Funding Acquisition).

## Data availability

Data supporting the conclusions of this article, along with scripts used for analysis and figure generation, are provided as supplemental materials or are available in the Coon laboratory’s GitHub repository (https://github.com/kcoonlab/stable-fly-AMR).

## Supplementary data

Supplemental File 1: Tables S1-S9

Supplemental File 2: PCR amplification of select ARGs

## References

1. Albuquerque TA, Zurek L. Temporal changes in the bacterial community of animal feces and their correlation with stable fly oviposition, larval development, and adult fitness. Front Microbiol 2014;5, DOI: 10.3389/fmicb.2014.00590.

2. Altschul SF, Gish W, Miller W et al. Basic local alignment search tool. Journal of Molecular Biology 1990;215:403–10.

3. Bagley ST, Seidler RJ. Primary Klebsiella identification with MacConkey-inositol-carbenicillin agar. Appl Environ Microbiol 1978;36:536–8.

4. Baldacchino F, Muenworn V, Desquesnes M et al. Transmission of pathogens by Stomoxys flies (Diptera, Muscidae): a review. Parasite 2013;20:26.

5. Becker K, Heilmann C, Peters G. Coagulase-Negative Staphylococci. Clin Microbiol Rev 2014;27:870–926.

6. Broce AB. An Improved Alsynite Trap for Stable Flies, Stomoxys calcitrans (Diptera: Muscidae). Journal of Medical Entomology 1988;25:406–9.

7. Butler JF, Kloft WJ, DuBose LA et al. Recontamination of Food After Feeding a 32P Food Source to Biting Muscidae. Journal of Medical Entomology 1977;13:567–71.

8. Castanheira M, Simner PJ, Bradford PA. Extended-spectrum β -lactamases: an update on their characteristics, epidemiology and detection. JAC-Antimicrobial Resistance 2021;3:dlab092.

9. CLSI. Performance Standards for Antimicrobial Susceptibility Testing; Twenty-Fifth Informational Supplement. CLSI Document M100-S25. Clinical and Laboratory Standards Institute, 2015.

10. CLSI. Clinical and Laboratory Standards Institute (CLSI). Performance Standards for Antimicrobial Susceptibility Testing. 34th Ed. CLSI Supplement M100. Clinical and Laboratory Standards Institute, 2024a.

11. CLSI. Performance Standards for Antimicrobial Disk and Dilution Susceptibility Tests for Bacteria Isolated From Animals. 7th Ed. CLSI Supplement VET01S. Clinical and Laboratory Standards Institute, 2024b.

12. Datsenko KA, Wanner BL. One-step inactivation of chromosomal genes in Escherichia coli K-12 using PCR products. Proc Natl Acad Sci USA 2000;97:6640–5.

13. Dougherty CT, Knapp FW, Burrus PB et al. Stable flies (Stomoxys calcitrans L.) and the behavior of grazing beef cattle. Applied Animal Behaviour Science 1993;35:215–33.

14. Feßler AT, Wang Y, Wu C et al. Mobile lincosamide resistance genes in staphylococci. Plasmid 2018;99:22–31.

15. Fournier C, Nordmann P, De La Rosa J-MO et al. KSA-1, a naturally occurring Ambler class A extended spectrum β-lactamase from the enterobacterial species Kosakonia sacchari. Journal of Global Antimicrobial Resistance 2024;39:6–11.

16. Frank JA, Reich CI, Sharma S et al. Critical Evaluation of Two Primers Commonly Used for Amplification of Bacterial 16S rRNA Genes. Appl Environ Microbiol 2008;74:2461–70.

17. Gelalcha BD, Kerro Dego O. Extended-Spectrum Beta-Lactamases Producing Enterobacteriaceae in the USA Dairy Cattle Farms and Implications for Public Health. Antibiotics 2022;11:1313.

18. Gonçalves JL, De Campos JL, Steinberger AJ et al. Incidence and Treatments of Bovine Mastitis and Other Diseases on 37 Dairy Farms in Wisconsin. Pathogens 2022;11:1282.

19. Goncalves JL, Mani R, Sreevatsan S et al. Apparent prevalence and selected risk factors of methicillin-resistant Staphylococcus aureus and non-aureus staphylococci and mammaliicocci in bulk tank milk of dairy herds in Indiana, Ohio, and Michigan. JDS Communications 2023;4:489– 95.

20. Hansens EJ. Fly Populations in Dairy Barns. Journal of Economic Entomology 1963;56:842–4.

21. Huttner A, Harbarth S, Carlet J et al. Antimicrobial resistance: a global view from the 2013 World Healthcare-Associated Infections Forum. Antimicrob Resist Infect Control 2013;2:31.

22. Karp BE, Tate H, Plumblee JR et al. National Antimicrobial Resistance Monitoring System: Two Decades of Advancing Public Health Through Integrated Surveillance of Antimicrobial Resistance. Foodborne Pathogens and Disease 2017;14:545–57.

23. Kim S-J, Moon DC, Park S-C et al. Antimicrobial resistance and genetic characterization of coagulase-negative staphylococci from bovine mastitis milk samples in Korea. Journal of Dairy Science 2019;102:11439–48.

24. Kolar QK, Goncalves JL, Erskine RJ et al. Comparison of Minimum Inhibitory Concentrations of Selected Antimicrobials for Non-Aureus Staphylococci, Enterococci, Lactococci, and Streptococci Isolated from Milk Samples of Cows with Clinical Mastitis. Antibiotics 2024;13:91.

25. Leininger DJ, Roberson JR, Elvinger F. Use of Eosin Methylene Blue Agar to Differentiate Escherichia Coli from Other Gram-Negative Mastitis Pathogens. J VET Diagn Invest 2001;13:273–5.

26. Lin X, Kück U. Cephalosporins as key lead generation beta-lactam antibiotics. Appl Microbiol Biotechnol 2022;106:8007–20.

27. Lüthje P, Von Köckritz-Blickwede M, Schwarz S. Identification and characterization of nine novel types of small staphylococcal plasmids carrying the lincosamide nucleotidyltransferase gene lnu(A). Journal of Antimicrobial Chemotherapy 2007;59:600–6.

28. MacAlpine JF ed. Manual of Nearctic Diptera. Vol. 2. Repr. Hull, Que: Canadian Gornvment Publ. Centre, 1993.

29. Makovec JA, Ruegg DrPL. Antimicrobial resistance of bacteria isolated from dairy cow milk samples submitted for bacterial culture: 8,905 samples (1994–2001). javma 2003;222:1582–9.

30. Martineau F, Picard FJ, Ke D et al. Development of a PCR Assay for Identification of Staphylococci at Genus and Species Levels. J Clin Microbiol 2001;39:2541–7.

31. McKenzie GJ, Craig NL. Fast, easy and efficient: site-specific insertion of transgenes into Enterobacterial chromosomes using Tn7 without need for selection of the insertion event. BMC Microbiol 2006;6:39.

32. Meyer JA, Petersen JJ. Characterization and Seasonal Distribution of Breeding Sites of Stable Flies and House Flies (Diptera: Muscidae) on Eastern Nebraska Feedlots and Dairies1. Journal of Economic Entomology 1983;76:103–8.

33. Murray CJL, Ikuta KS, Sharara F et al. Global burden of bacterial antimicrobial resistance in 2019: a systematic analysis. The Lancet 2022;399:629–55.

34. Ogutu JO, Zhang Q, Huang Y et al. Development of a multiplex PCR system and its application in detection of blaSHV, blaTEM, blaCTX-M-1, blaCTX-M-9 and blaOXA-1 group genes in clinical Klebsiella pneumoniae and Escherichia coli strains. J Antibiot 2015;68:725–33.

35. Oliver SP, Murinda SE, Jayarao BhushanM. Impact of Antibiotic Use in Adult Dairy Cows on Antimicrobial Resistance of Veterinary and Human Pathogens: A Comprehensive Review. Foodborne Pathogens and Disease 2011;8:337–55.

36. Pagani L, Dell’Amico E, Migliavacca R et al. Multiple CTX-M-Type Extended-Spectrum β- Lactamases in Nosocomial Isolates of Enterobacteriaceae from a Hospital in Northern Italy. J Clin Microbiol 2003;41:4264–9.

37. Patel SJ, Wellington M, Shah RM et al. Antibiotic Stewardship in Food-producing Animals: Challenges, Progress, and Opportunities. Clinical Therapeutics 2020;42:1649–58.

38. Pfeifer Y, Cullik A, Witte W. Resistance to cephalosporins and carbapenems in Gram-negative bacterial pathogens. International Journal of Medical Microbiology 2010;300:371–9.

39. Redding LE, Bender J, Baker L. Quantification of antibiotic use on dairy farms in Pennsylvania. Journal of Dairy Science 2019;102:1494–507.

40. Ruegg PL. Realities, Challenges and Benefits of Antimicrobial Stewardship in Dairy Practice in the United States. Microorganisms 2022;10:1626.

41. Sasaki T, Kobayashi M, Agui N. Epidemiological Potential of Excretion and Regurgitation by Musca domestica (Diptera: Muscidae) in the Dissemination of Escherichia coli O157: H7 to Food. me 2000;37:945–9.

42. Sawant AA, Sordillo LM, Jayarao BM. A Survey on Antibiotic Usage in Dairy Herds in Pennsylvania. Journal of Dairy Science 2005;88:2991–9.

43. Schlechter RO, Jun H, Bernach M et al. Chromatic Bacteria – A Broad Host-Range Plasmid and Chromosomal Insertion Toolbox for Fluorescent Protein Expression in Bacteria. Front Microbiol 2018;9:3052.

44. Schoenfelder SMK, Dong Y, Feßler AT et al. Antibiotic resistance profiles of coagulase-negative staphylococci in livestock environments. Veterinary Microbiology 2017;200:79–87.

45. Showler AT, Osbrink WLA. Stable Fly, Stomoxys calcitrans (L.), Dispersal and Governing Factors. Int J Insect Sci 2015;7:IJIS.S21647.

46. Sommer AJ, Deblois CL, Tu ADJ et al. Opportunistic pathogens are prevalent across the culturable exogenous and endogenous microbiota of stable flies captured at a dairy facility. bioRxiv 2024 Nov 4;10.1101/2024.11.04.621909 [Preprint].

47. Sommer AJ, Kettner JE, Coon KL. Stable flies are bona fide carriers of mastitis-associated bacteria. mSphere 2024;9:e00336–24.

48. Stelder JJ, Kjær LJ, Jensen LB et al. Livestock-associated MRSA survival on house flies (Musca domestica) and stable flies (Stomoxys calcitrans) after removal from a Danish pig farm. Sci Rep 2021;11:3527.

49. Taponen S, Pyorala S. Coagulase-negative staphylococci as cause of bovine mastitis—Not so different from Staphylococcus aureus? Veterinary Microbiology 2009;134:29–36.

50. Thornsberry C, Marler JK, Watts JL et al. Activity of pirlimycin against pathogens from cows with mastitis and recommendations for disk diffusion tests. Antimicrob Agents Chemother 1993;37:1122–6.

51. Weill F-X, Demartin M, Tandé D et al. SHV-12-Like Extended-Spectrum-β-Lactamase-Producing Strains of Salmonella enterica Serotypes Babelsberg and Enteritidis Isolated in France among Infants Adopted from Mali. J Clin Microbiol 2004;42:2432–7.

52. Wendlandt S, Kadlec K, Feßler AT et al. Identification of ABC transporter genes conferring combined pleuromutilin–lincosamide–streptogramin A resistance in bovine methicillin-resistant Staphylococcus aureus and coagulase-negative staphylococci. Veterinary Microbiology 2015;177:353–8.

53. Zhao S, White DG, McDermott PF et al. Identification and Expression of Cephamycinase bla CMY Genes in Escherichia coli and Salmonella Isolates from Food Animals and Ground Meat. Antimicrob Agents Chemother 2001;45:3647–50.

