## Supplementary material for "Prevalence of antimicrobial resistance phenotypes and genes in stable fly- and manure-derived bacterial isolates from clinically relevant taxa in dairy settings": File S2

**File S2**. PCR amplification of select ARGs.

*----lnuA* PCR----

Row 1 wells (left to right):

100BP ML, AS0204, AS0295, AS0300, AS0307, AS0308, AS0311, AS0554, AS0569, AS0602, AS0650, AS0913

Row 2 wells (left to right):

100BP ML, AS1275, AS1301, AS1397, AS1400, AS1404, AS1411, AS1442, AS1445, molecular water

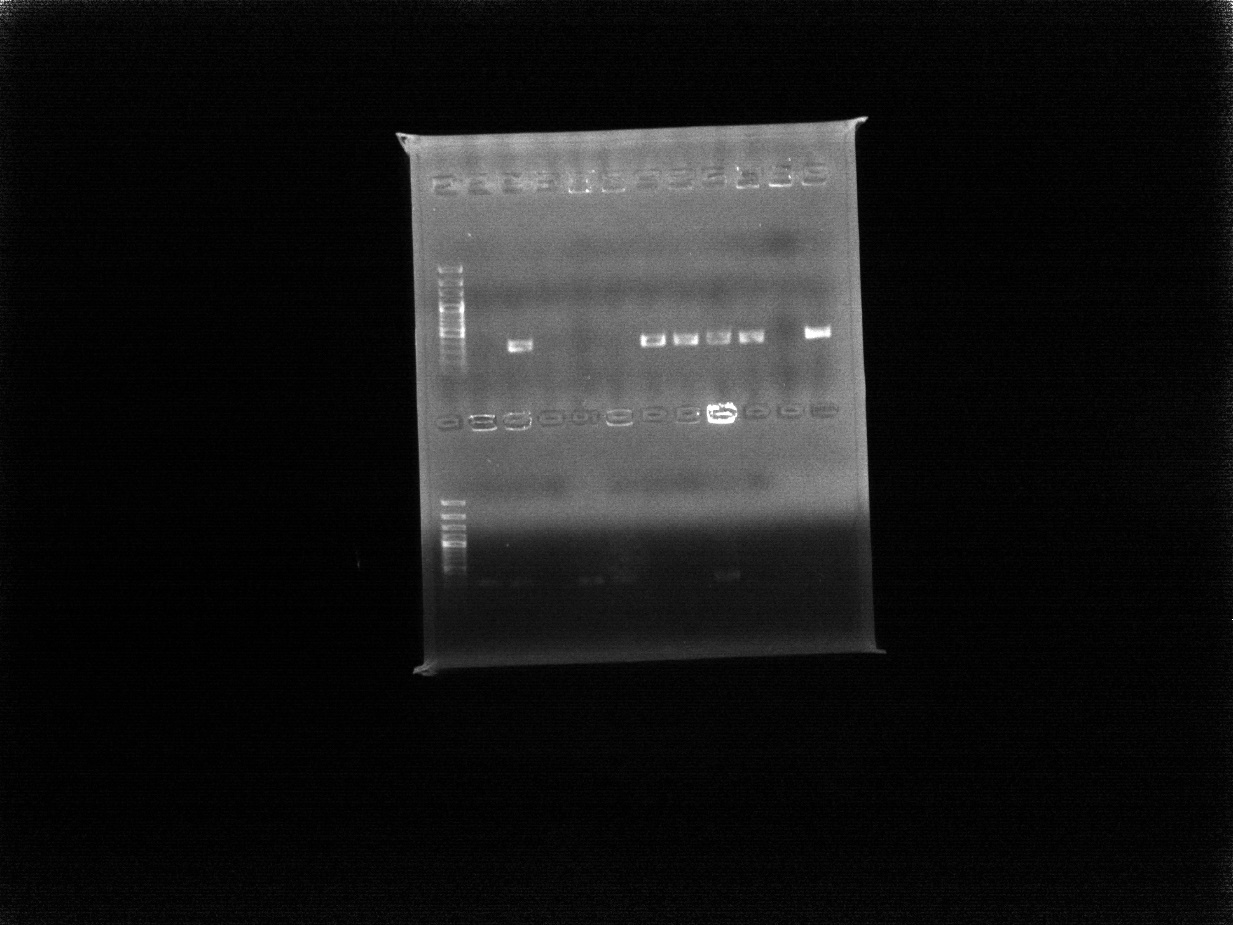

*lnuA* positive (323bp band):

AS0295, AS0311, AS0554, AS0569, AS0602, AS0913, AS1275, AS1301, AS1400, AS1404, AS1445

*----bla*OXA PCR----

Row 1 wells (left to right):

100BP ML, AS0233, AS0234, AS0466, AS0507, AS0468, AS0690, AS0540, AS1210

Row 2 wells (left to right):

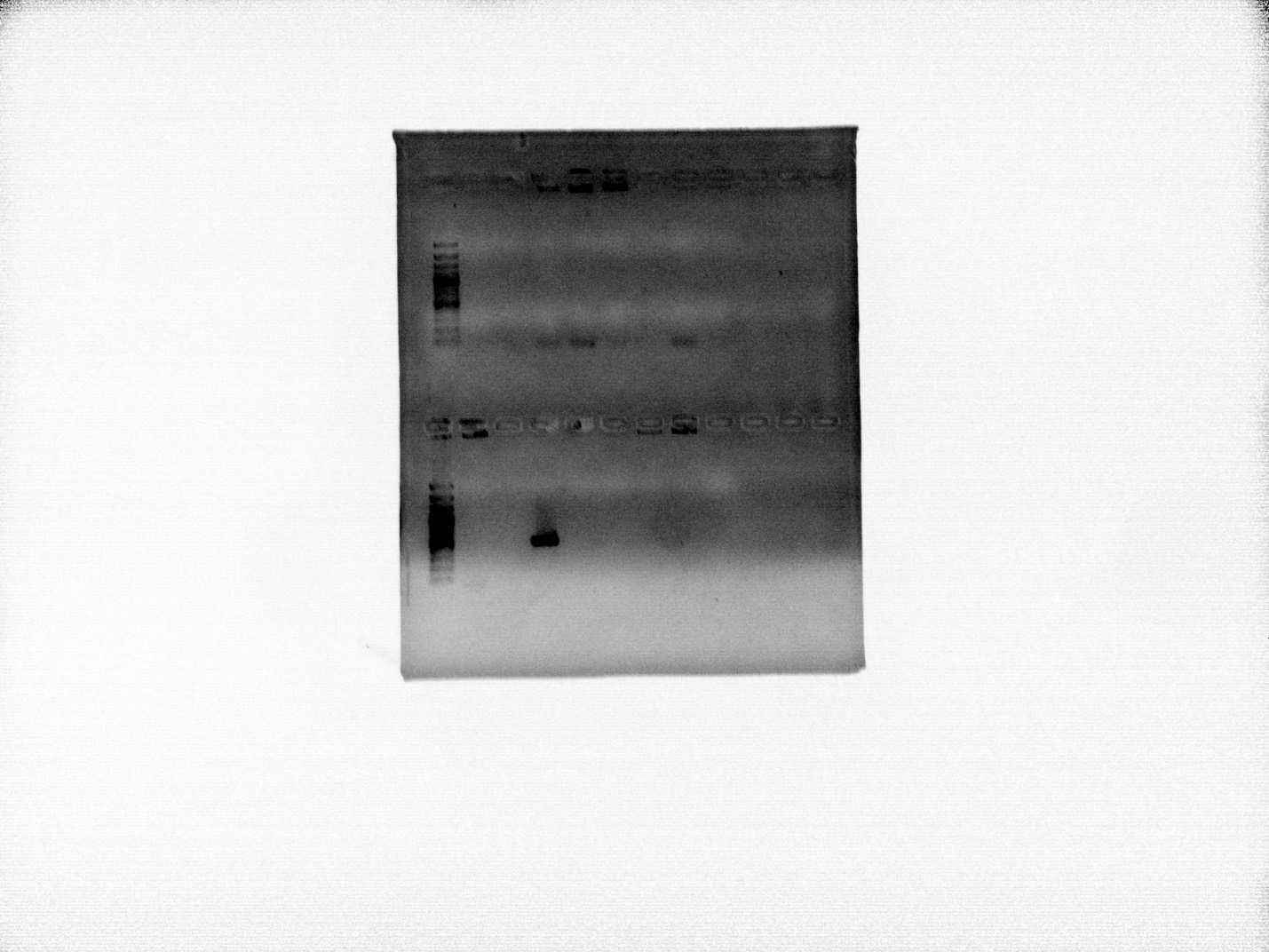
100BP ML, AS1211, AS0255, AS0467, AS1245, AS1257, AS1151, AS1195, molecular water

*lnuA* positive (564bp band):

AS0467

*----bla*CTX PCR----

Row 1 wells (left to right):

100BP ML, AS0233, AS0234, AS0466, AS0507, AS0468, AS0690, AS0540, AS1210

Row 2 wells (left to right):

100BP ML, AS1211, AS0255, AS0467, AS1245, AS1257, AS1151, AS1195, molecular water

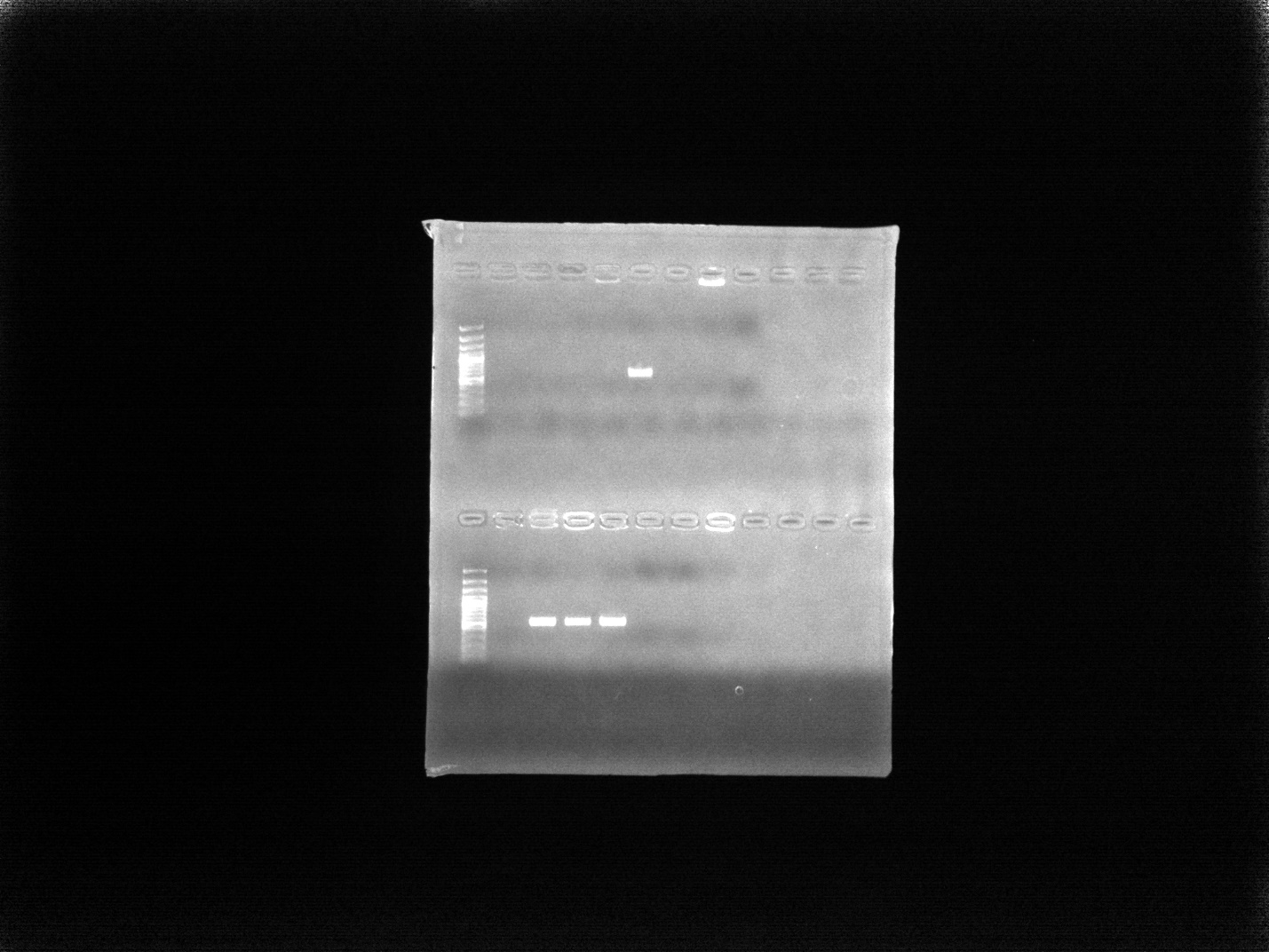

CTX positive (593bp band):

AS0468, AS0255, AS0467, AS1245

*----bla*TEM PCR----

Row 1 wells (left to right):

100BP ML, AS0233, AS0234, AS0466, AS0507, AS0468, AS0690, AS0540, AS1210

Row 2 wells (left to right):

100BP ML, AS1211, AS0255, AS0467, AS1245, AS1257, AS1151, AS1195, molecular water

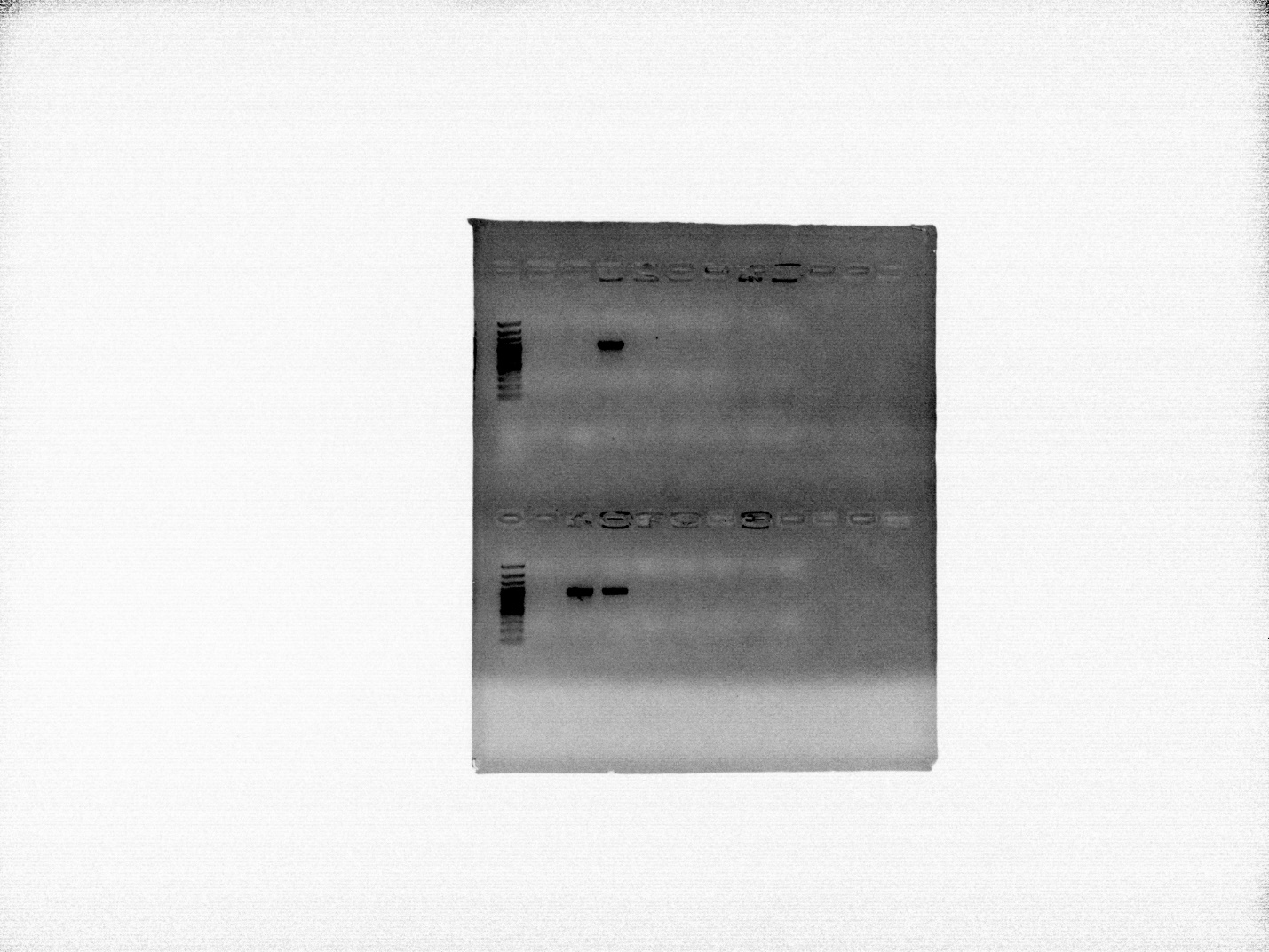

TEM positive (1086bp band):

AS0466, AS0255, AS0467

*----bla*CMY PCR----

Row 1 wells (left to right):

100BP ML, AS0233, AS0234, AS0466, AS0507, AS0468, AS0690, AS0540, AS1210

Row 2 wells (left to right):

100BP ML, AS1211, AS0255, AS0467, AS1245, AS1257, AS1151, AS1195, molecular water

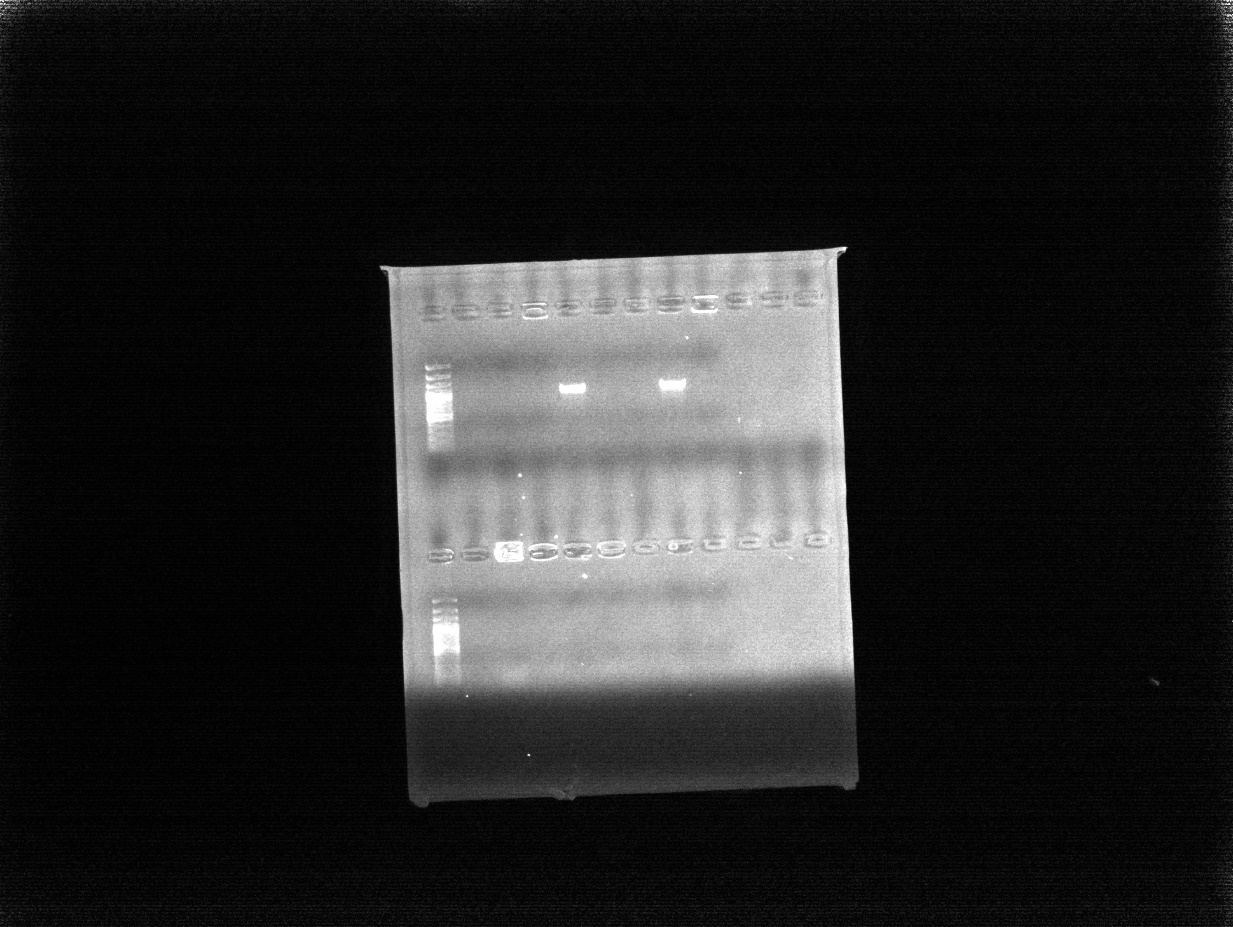
CMY positive (1000bp band):

AS0507, AS0540
